# Distinct core promoter codes drive transcription initiation at key developmental transitions in a marine chordate

**DOI:** 10.1101/140764

**Authors:** Gemma B. Danks, Pavla Navratilova, Boris Lenhard, Eric Thompson

## Abstract

Development is largely driven by transitions between transcriptional programs. The initiation of transcription at appropriate sites in the genome is a key component of this and yet few rules governing selection are known. Here, we used cap analysis of gene expression (CAGE) to generate bp-resolution maps of transcription start sites (TSSs) across the genome of *Oikopleura dioica*, a member of the closest living relatives to vertebrates. Our TSS maps revealed promoter features in common with vertebrates, as well as striking differences, and uncovered key roles for core promoter elements in the regulation of development. During spermatogenesis there is a genome-wide shift in mode of transcription initiation characterized by a novel core promoter element. This element was associated with > 70% of transcription in the testis, including the male-specific use of cryptic internal promoters within operons. In many cases this led to the exclusion of *trans*-splice sites, revealing a novel mechanism for regulating which mRNAs receive the spliced leader. During oogenesis the cell cycle regulator, E2F1, has been co-opted in regulating maternal transcription in endocycling nurse nuclei. In addition, maternal promoters lack the TATA-like element found in vertebrates and have broad, rather than sharp, architectures with ordered nucleosomes. Promoters of ribosomal protein genes lack the highly conserved TCT initiator. We also report an association between DNA methylation on transcribed gene bodies and the TATA-box, which indicates that this ancient promoter motif may play a role in selecting DNA for transcription-associated methylation in invertebrate genomes.

## Introduction

Sites for the initiation of transcription are frequently marked in the genome by specific sequence elements, which are recognized and subsequently bound by basal transcription factors [1, 2]. The diversity of core promoter elements suggests that they play important roles in the differential regulation of subsets of genes. For example, the conserved TATA-box, which is bound by TATA-binding protein, is responsible for transcription initiation at tissue-specific promoters in metazoans [3], whereas a degenerate TATA-like element is associated with maternal transcription initiation in vertebrates [4]. Other core elements may be critical to development, but as yet none has been assigned a specific role(s).

In vertebrates, the role of 5-methylcytosine DNA methylation is location-dependent: it is associated with repression when located at promoters and active transcription and splicing when located within gene bodies [5, 6]. DNA methylation within active gene bodies is a feature that is conserved between animals and plants (although lost in certain lineages including *Caenorhabditis elegans* and *Drosophila*) [5] and represents the majority of DNA methylation in the genome of the urochordate, ascidian, *Ciona intestinalis* [7, 8]. One function of DNA methylation in gene bodies is repression of alternative intragenic promoters [9, 10]. In invertebrate genomes, gene body methylation is only found at a subset of genes and is positively correlated to gene expression level [7, 8], but how this subset is selected for methylation has so far remained unknown.

The identification of core promoter elements, and mapping of transcription start sites (TSSs), at singlenucleotide resolution, has been facilitated by Cap Analysis of Gene Expression (CAGE) [11]. This has led to the discovery of two main modes for specifying sites of transcription initiation [2, 12]. Sequence motifs bound by the pre-initiation complex result in transcription initiation within a narrow region and lead to “sharp” promoter architectures. Conversely, the positioning of nucleosomes defines a wider catchment area for the pre-initiation complex and leads to “broad” promoter architectures [4]. Promoter architectures can also show associations to downstream translational events. For example, promoters of ribosomal protein genes are usually sharp with a highly conserved TCT Initiator (Inr) sequence [2, 13, 14], which forms the beginning of a Terminal OligoPyrimidine (TOP) motif critical for nutrient-dependent translational control [15]. In mammals, these promoters, unusually, have both a TATA-box and CpG islands. In *C. intestinalis* they are sharp with a TCT initiator, but lack a TATA-box [13]. Recently, it has been shown that a genome-wide switch occurs in the mode of TSS selection during vertebrate embryogenesis [4]. Maternal promoters in vertebrates are sharp, or multiple sharp, with TATA-like, AT-rich (W-box) upstream elements guiding TSS selection. During the maternal to zygotic transition, nucleosomes with H3K4me3 are positioned at zygotic promoters that lack a W-box, leading to broad promoter architectures. The extent to which these, or similar, features are evolutionarily conserved is unknown.

*Oikopleura dioica* is a marine, larvacean, chordate in the sister group to vertebrates and is well positioned to examine the evolution of TSS features and the dynamics of TSS selection. The *O. dioica* genome is the most compact of any animal genome sequenced so far and 28% of its genes are organised into operons [16]. Each operon contains two or more genes that are transcribed from a single promoter located upstream of the first gene. The resulting polycistronic mRNA is resolved via the *trans*-splicing [17, 18] of a spliced-leader (SL) sequence to unpaired acceptor sites at the 5’ ends of each resulting monocistron. *Trans*-splicing in *O. dioica* [19] also occurs at monocistronic genes; 39% of all annotated genes give rise to mRNAs that are *trans*-spliced [20]. During *trans*-splicing a portion of the original 5’ sequence upstream of the *trans*-splicing acceptor site is removed. Here, we mapped TSSs at single-nucleotide resolution, using CAGE, in six key stages of *O. dioica*, development, covering the entire 6-day life cycle. In order to maximise the mapping of original TSSs (rather than trans-splice sites) we sequenced only mRNAs without the SL sequence. We used our TSS maps, together with previously generated genome-wide maps of *trans*-splice sites [20], E2F1 binding sites, key histone modifications and DNA methylation [21], to derive TSS-selection criteria at major developmental transitions. Our data show that *O. dioica* has some promoter features in common with vertebrates, including evidence of nucleosome positioning at broad promoters and tissue-specific expression of TATA-dependent promoters, but it differs markedly in its mode of maternal transcription initiation, which is characterized by the ordering of nucleosomes and the binding of the cell cycle regulator E2F1. *O. dioica* also employs a remarkable genome-wide shift in mode of TSS-selection during spermatogenesis, associated with a distinct, tissue-specific, TCTAGA core promoter motif, that has not been previously identified.

## Results

### Promoter usage across development

We extracted RNA for CAGE from *O. dioica* at six stages of development across the 6-day life cycle of the animal (Figure 1A). Illumina sequencing generated >39 M reads, of which 2.4–5.9 M (54–64%) for each stage mapped uniquely to the genome (Table S1). Summing tags that mapped to unique positions gave the abundance of transcripts originating from each TSS. We normalized these counts to tags per million reads (tpm) and clustered neighbouring TSSs to generate tag clusters (TCs), which revealed the set of promoter regions that are active within each stage. TCs (supported by at least 1 tpm in at least one stage) mapped to 6,241 annotated genes, 4,937 of which were defined as expressed using previously generated tiling array data [22] across equivalent developmental stages (Figure S1). Multiple genes within an operon are transcribed from a common TSS. In line with this we captured TCs for only 538 downstream operon genes, out of 2,832 (19%) that were defined as expressed based on tiling array data (Figure S1). TCs for these 538 genes include previously unidentified stage-specific use of cryptic internal promoters within operons. Previously, we generated a bp-resolution, genome-wide map of *trans*-splice sites in *O. dioica* [20] using pooled animals collected at the same developmental stages we used here. As previously, we define a gene as *trans*-spliced if it is associated with a mapped *trans*-splice site. Our newly generated CAGE dataset captures the original TSSs of 51% (1341/2643) of all monocistronic (non-operon) *trans*-spliced genes allowing us to analyse promoter features of *trans*-spliced genes.

**Figure 1:**
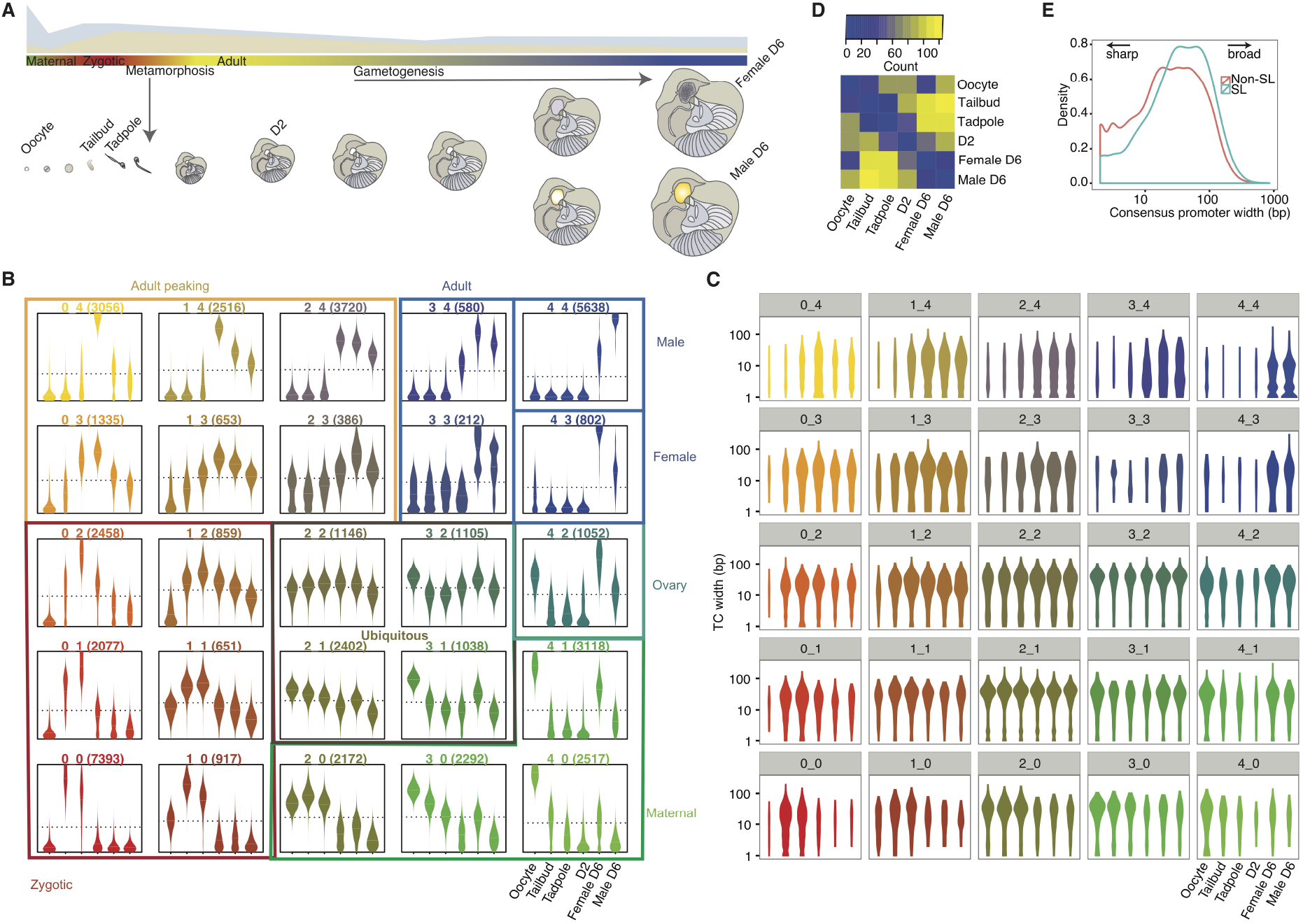
Promoter usage across the *O. dioica* life cycle. (A) The 6-day life cycle with stages used for singlenucleotide resolution of TSS-mapping by CAGE are labelled (oocyte, tailbud, tadpole, day 2, female day 6 and male day 6). Changes in the proportion of mRNAs that are *trans*-spliced (blue) and non-*trans*-spliced (beige) are shown schematically in the upper panel, above a colour bar indicating major promoter categories (colours corresponding to clusters in (B)). Developmental stages are shown schematically below this bar. (B) Expression profiles obtained from self-organising map clustering of CAGE TSSs (CTSSs). Each beanplot shows the distribution of relative expression (y-axis) originating at CTSSs (number of CTSSs above each plot) within each cluster at each developmental stage (x-axis) labelled only in the bottom right plot. Coloured boxes and associated labels indicate groups of clusters with similar expression profiles. (C) Beanplots showing the distribution of interquantile widths of tag clusters (TCs) within each stage and assigned to the expression cluster of the dominant CTSS (plots are ordered and coloured as in (B) revealing an increase in the use of sharp promoters in adult(tissue)-specific genes. (D) Heatmap showing the number of promoters that shift up- or down-stream in location between all possible pairs of developmental stages. The highest number of shifting promoters occurs between pre-metamorphic (tailbud) and post-metamorphic (day 2 and day 6) stages. (E) Distribution of the interquartile widths of consensus promoter regions of *trans*-spliced (SL) and non-*trans*-spliced (Non-SL) genes.

We defined 13,771 consensus promoter regions in the genome by clustering TCs, with > 5 tpm, across stages [23, 4]. Expression profiles of individual TSSs were clustered using a self-organizing map [23, 4] (SOM) in order to assess the dynamics of TSS selection across development (Figure 1B). Distinct ubiquitous, maternal and zygotic expression TSS clusters were present as well as a large cluster of male-specific TSSs, related to transcription initiation sites associated with spermatogenesis. SOM clustering of consensus promoter region expression profiles revealed similar patterns (Figure S2).

### A genome-wide shift in mode of TSS selection during spermatogenesis

Maternal and ubiquitously expressed TSSs (identified by SOM clustering; Figure 1B), and TSSs associated with *trans*-spliced genes, were found predominantly within broad TCs (Figure 1C,E and Figure S2) whereas TSSs used in adult stages, particularly male-specific (hereafter referred to as spermatogenic) TSSs, were predominantly found in sharp TCs (Figure 1C, Figure S2). The presence of sharp TCs suggests sequence motifs in these core promoters determine the selection of TSSs at a fixed distance downstream. We therefore examined all promoter sequences and identified a core promoter element (TCTAGA), embedded in a TT-rich sequence context, which was remarkably specific to spermatogenic TSSs (Figure 2 and Figure S3). This element was present in 71.6% (1391/1943) of spermatogenic TCs in the male and was strictly positioned 40–50 nt upstream of the dominant TSS (with a strong preference for 45–48 bp; Figure 2B), suggesting the binding of a male-specific variant of the TFIID complex, components of which are known to play roles in development and gametogenesis across metazoans [24]. We re-analyzed existing CAGE data [23, 25] from a time course of 8 testis samples across mouse development from embryogenesis to adult tissues. We found no enrichment for a position-specific TCTAGA motif in promoter regions of any stage, nor of promoters with spermatogenesis-associated expression patterns (data not shown). We searched the promoter regions of 16,671 annotated genes in the *C. intestinalis* genome and found only 226 (1.3%) with a TCTAGA within 100 bp upstream of the annotated start site compared to 2088 (12.5%) that had a consensus TATA-box motif. This suggests a larvacean, lineage-specific evolution of this mode of TSS-selection for the activation of the spermatogenesis transcriptional program.

**Figure 2:**
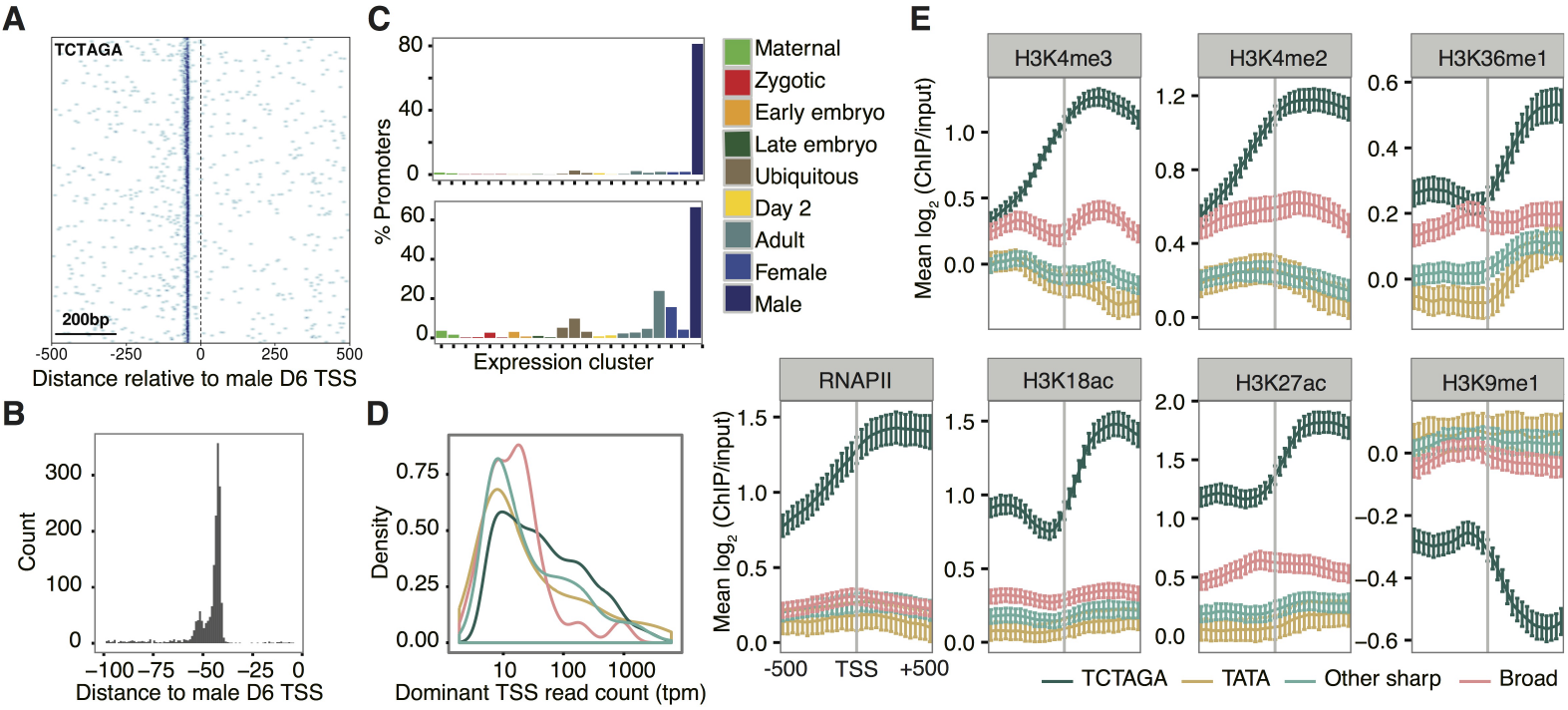
Features of spermatogenic promoters in *O. dioica*. (A) Heatmap shows the density of TCTAGA at each position (x-axis) in a −500 to +500 bp region centred on the male dominant TSS for spermatogenic promoter sequences (rows) ordered by promoter width (top to bottom = broad to sharp). Darker blue indicates higher enrichment. (B) Distribution of distances (median = 44 bp) to the TCTAGA motif relative to the dominant TSS in male day 6 animals. (C) The TCTAGA motif is specific to spermatogenic transcription: of all promoters with a TCTAGA motif the majority have a male-specific expression profile (top). Moreover, the majority of spermatogenic transcription is associated with a TCTAGA motif: of all promoters with a male-specific expression profile the majority contain a TCTAGA, whereas very few promoters in other expression classes contain this motif (bottom). (D) Distribution of tag counts (tpm) for different promoter classes in the male (TATA = TSS with upstream TATAW element; TCTAGA = TSS with upstream TCTAGA; Sharp = all other promoters with narrow region of TSSs; Broad = dispersed region of TSSs). (E) ChIP-chip data for day 6 testes are shown for RNAPII, H3K18ac, H3K27ac, H3K4me3, H3K4me2, H3K36me1 and H3K9me1. Each plot shows the mean log_2_ ratio of ChIP/input at each probe position in a 1000 bp window centered on the dominant TSS. Promoters were categorized according to promoter type. Error bars show 95% confidence intervals for the mean obtained by bootstrapping.

We next identified promoter regions with a shift in TSS usage [23, 4] between any pair of developmental stages. When comparing the embryo versus adult male stages (Figure 1D) we observed the highest number (124/519) of single promoter regions for which the TSS location changed (< 40% TC overlap). In 43/124 of these cases there was a shift from a TCTAGA-independent promoter in the embryo to a TCTAGA-associated promoter in the male.

We identified 693 cryptic internal promoters [26] within operons: spermatogenic promoters (208) were over-represented and the TCTAGA element was found more frequently (25.5%; 177 promoters) than expected (*χ*^2^ = 142.98, df = 1, *p* < 2.2 × 10^−16^). We then analyzed patterns of enrichment of the H3K4me3 promoter mark from ChIP-chip data [21] in the ovary and testis of animals at the same developmental stage as the day 6 male and female animals used to generate our CAGE dataset. In support of the presence of spermatogenic-specific cryptic internal promoters within operons, we only found enrichment of H3K4me3 at the start sites of internal genes within operons in the testis, whereas the start sites of operons were enriched for this mark in both the ovary and testis (Figure S4). We performed 5’ RACE for 12 spermatogenic, TCTAGA TSSs that were located in operons in front of internal cistron units that were expressed in both the testis and ovary (expression determined using microarray data [22]). In 5/12 cases we found only the male-specific TSS in males, with no evidence of the SL sequence, and only the *trans*-spliced form in 4/5 of these in females. This indicates that the associated gene is transcribed as part of the expected polycistron in females but predominantly as non-*trans*-spliced monocistrons in males from male-specific cryptic internal promoters. In 1/12 cases we did not detect the male-specific TSS or a *trans*-spliced form in either male or females. In 6/12 cases we detected only *trans*-spliced 5’ ends in males and females as well as testes samples. In 5/6 of these the *trans*-splice site was upstream of the male-specific CAGE TSS, indicating that these genes were predominantly found as part of polycistronic transcripts and that the TCTAGA promoter was used at a lower frequency.

Sites for *trans*-splicing are determined by the presence of an unpaired AG acceptor site, which is usually followed by an adenine [20]. Remarkably, we found that 89 spermatogenic promoters in males (associated with 87 genes) had a TCTAGA motif with its AGA mapping to a *trans*-splice site (representing 16.4% of all TCTAGA spermatogenic genes that were annotated as *trans*-spliced). Transcription downstream of these TCTAGA elements during spermatogenesis therefore results in mRNAs that lack a *trans*-splice acceptor site and are therefore not *trans*-spliced with the SL sequence. Transcription driven by alternative upstream promoters during other stages of development leads to mRNAs with the *trans*-splice site intact and are therefore *trans*-spliced with the SL. This finding reveals a novel mechanism for the developmental regulation of *trans*-splicing.

Spermatogenic TCTAGA promoters had significantly higher expression levels compared to other promoter types in males (all *p* < 0.05; Figure 2D). We analyzed the profiles of a range of histone modifications as well as RNA pol II occupancy using ChIP-chip in the testis and ovaries of day 6 stage-matched animals [21]. We found that spermatogenic TCTAGA promoters were associated with higher RNA pol II occupancy and higher enrichment of histone modifications associated with active transcription (and depletion of repressive marks) in the testis, including specific marking by H3K18ac (Figure 2E). Several of these marks were independent of expression level (Figure S5). Together, our data revealed a unique transcription initiation code that was specific to spermatogenic core promoters. This code is associated with a chromatin state primed for high levels of transcription in the testis and directs both a genome-wide shift in promoter usage, and the developmental regulation of operon transcription and *trans*-splicing.

### Ordered nucleosomes and E2F1 determine maternal TSS-selection in endocycling nurse nuclei

Maternal promoters in vertebrates tend to be sharp, or multiple sharp, with a degenerate TATA-like motif (W-box) determining TSSs [4]. In contrast, we found that maternal promoters in *O. dioica* were broad (Figure 1C) and lacked a W-box at the expected TATA-box position or any other enrichment of dinucleotides (Figure S3). Broad promoters in vertebrates are associated with ordered nucleosomes, as shown by the precise positioning of histone H3K4me3 enrichment at the first nucleosome downstream from the dominant TSS [4]. Here, we used ChIP-chip data [21] from the ovaries of day 6 (stage-matched) *O. dioica* and analyzed the profiles of H3 and H3K4 histone modifications around dominant TSSs of maternal promoters. Distinct peaks of histone H3 enrichment flanked the dominant TSSs at broad promoters in the ovary (Figure 3A) with a peak in H3K4me3 enrichment immediately downstream (Figure 3A) as seen in vertebrate broad promoters. These data showed that TSS-selection in *O. dioica* broad promoters has similar features to those in vertebrate broad promoters and is the main mode of maternal TSS selection in *O. dioica*.

**Figure 3:**
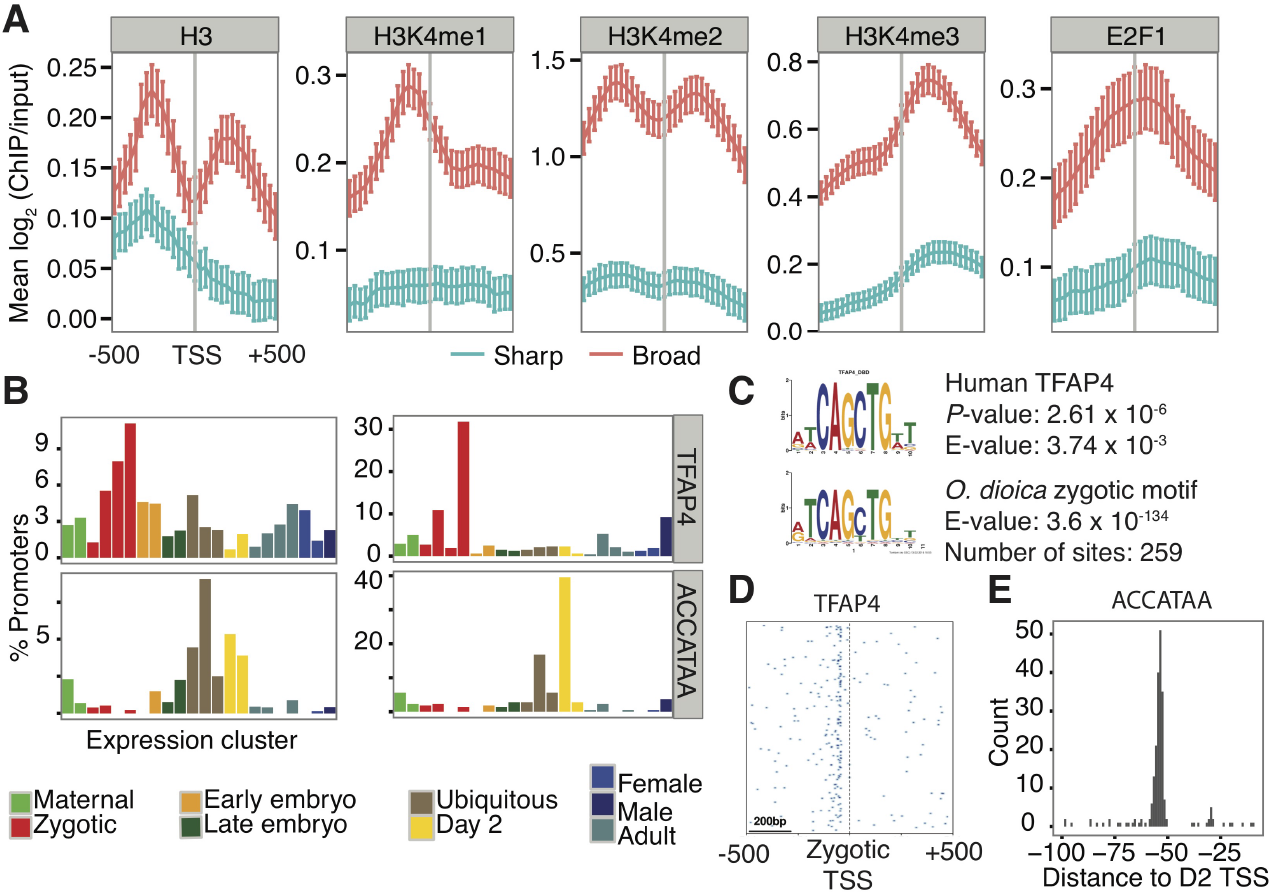
Features of maternal, zygotic and ubiquitous promoters in *O. dioica*. (A) Ordered nucleosome positioning at broad promoters in the ovary. Data are shown for H3, H3K4me1, H3K4me2, H3K4me3 and E2F1 ChIP-chip experiments. Each plot shows the mean log_2_ ratio of ChIP/input at each probe position in a 1000 bp window centred on the dominant TSS. Promoters were categorized into sharp (blue) and broad (pink) using lower and upper quartiles of widths across all stages. Error bars show 95% confidence intervals for the mean obtained by bootstrapping. (B) Percentage of all consensus promoters (left) within each expression cluster (x-axis) (profiles shown in Figure S2A) containing sequence elements as labelled (*O. dioica* TFAP4-like motif, ACCATAA motif). Percentage of promoters that contain each motif falling within each expression cluster is also shown (right). Expression clusters are grouped and coloured as coded in Figure 1C. (C) Sequence logo for an over-represented motif (E-value and number of sites as indicated) in zygotic promoters (tailbud sequences from TCs with dominant CTSS in cluster 0_0) in *O. dioica* (top), and its alignment to a significant match (E-value and *p* as indicated) for the binding site motif of human TFAP4. (D) Heatmap shows the density of the zygotic promoter motif matching TFAP4 at each position (x-axis) in a −500 to +500 bp region centred on the tailbud dominant TSS for zygotic promoter sequences (rows) ordered by promoter width (top to bottom = broad to sharp). Darker blue indicates higher enrichment. (E) Distance relative to the dominant TSS in day 2 animals for the ACCATAA motif found in sharp promoters that have ubiquitous and day 2-specific expression profiles.

We found that the nucleosome-free region at the TSS corresponds to an enrichment of the activating transcription factor E2F1 (Figure 3A), a key regulator of the cell cycle [27]. A remarkable 27.7% (1075/3882) of genes with strong CAGE support (≥ 5 tpm) in the day 6 female had maternal promoters in the ovary bound by E2F1. These genes were enriched for, though not limited to, known E2F1-regulated functions (Figure S6). These results suggest that, in addition to its known roles in cell cycle-related regulation of transcription, there is a co-option of E2F1 as a regulator of maternal transcription in *O. dioica*.

Maternal promoters in *O. dioica* were located on the X-chromosome more frequently than expected, compared to zygotic promoters (*χ*^2^ = 43.34, df = 1, *p* = 4.61 × 10^−11^), and there was a corresponding enrichment of E2F1 on the X-chromosome in the ovary: 40% (634/1566) of female TCs on the X-chromosome were bound by E2F1 compared to 23% (975/4250) of female TCs on autosomes (*χ*^2^ = 179.9382, df = 2, *p* < 2.2 × 10^−16^). This enrichment was present even when considering only promoters with high CpG content, which are more common on the X-chromosome. These results revealed a female-bias of X-linked genes [28] in *O. dioica*.

### Regulation of zygotic promoters

Zygotic promoters in *O. dioica* (TSS clusters with low maternal and high embryonic expression; Figure 1B) contained an upstream GC-rich region, characteristic of broad promoters, and a downstream poly(T)-tract (Figure S3). An E-box [29] motif with a significant match to the binding site of TFAP4 (activating enhancer binding protein 4), a regulator of cell proliferation, was over-represented in the region immediately upstream of 259 zygotic-specific TSSs in the embryonic tailbud stage (Figure 3B-D). This suggests a TFAP4-like factor may be a key regulator in the tailbud, the stage critical for mitotic cell proliferation, differentiation and organogenesis.

### TSS selection in ubiquitous and ribosomal protein gene promoters

Most *O. dioica* promoters used to drive ubiquitous expression throughout the life cycle (Figure 1B) had a broad architecture with a strong GC-rich band immediately upstream of the TSS, as seen in zygotic promoters, and a clear GAAA signal at the expected +1 nucleosome position (Figure S3). We also found a position-specific (median distance 56 bp upstream; Figure 4E) ACCATAA sequence element associated with TSS-selection in sharp ubiquitous promoter regions (Figure 3C and Figure S3), as well as in sharp promoters specific to day 2 animals (juvenile animals; pre-gametogenesis). This motif was present in 215 consensus promoter regions.

**Figure 4:**
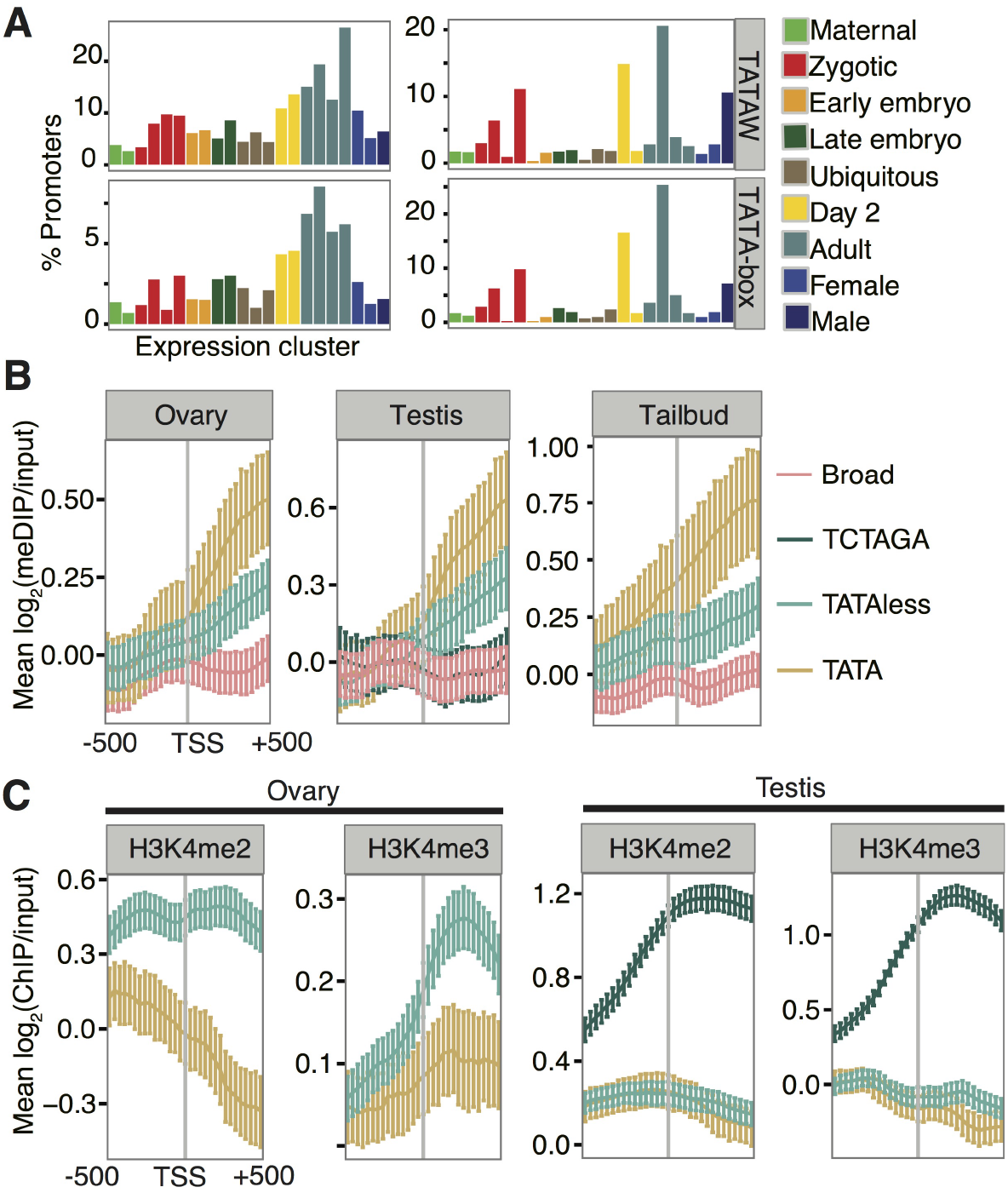
Features of TATA-dependent promoters in *O. dioica*. (A) Percentage of all consensus promoters within each expression cluster (left) (profiles shown in Figure S2A) containing TATA-elements as labelled (TATAW = core TATA motif; TATA-box = consensus TATA-box). Percentage of promoters that contain TATA-elements within each expression cluster is also shown (right): the majority of TATA-dependent promoters are adult-specific. Expression clusters are grouped and coloured according to colours in Figure 1C. B-C Plots show the mean log_2_ ratio of methyl-DNA IP/input (B) or ChIP/input (C) (y-axis) at each probe position (x-axis) in a 1000 bp window centred on the dominant TSS, in ovaries, testes and embryos (tailbud) as labelled. Error bars show 95% confidence intervals for the mean obtained by bootstrapping. Promoters were divided into broad and sharp with sharp subdivided into those with a TATAW-element and those without and, in the testis, those with the position-specific TCTAGA motif. DNA methylation was enriched in the downstream of TATA-dependent promoters (B) and H3K4me2 and H3K4me3 were depleted in the same regions (C).

Whereas a typical Initiator (Inr) CA dinucleotide was present in 53% of consensus promoter regions in *O. dioica*, the highly conserved TCT initiator was absent from all CAGE-detected ribosomal protein genes (29 detected out of 129 annotated; the majority being located within operons [20]). TCs of these genes were predominantly broad (only 6/51 were sharp; 2/6 contained a TATA-element), in line with other *trans*-spliced gene promoters (Figure 1D) with a higher average CpG content than non-ribosomal protein genes (Welch Two Sample t-test: t = 3.22, df = 35.164, *p* = 1.379 × 10^−3^). This indicates that these promoters have lost the specific transcriptional regulation conferred by the TCT initiator in other species and provides further evidence for the role of the SL in replacing the role of the TOP motif [20].

### Conserved tissue-specific TATA-dependent TSS-selection is associated with higher levels of DNA methylation in gene bodies

A core promoter TATAW element was present in 10.7% of consensus promoter regions, in line with the percentages of TATA-dependent promoters in mammals [30]. A lower percentage (3.8%) of promoters had a longer consensus TATA-box motif (TATAWAWR). The use of promoters with this consensus motif was specific to sharp promoters in adult stages (Figure 4A and Figure S3), indicating that this mode of TSS-selection at tissue-specific promoters is conserved between *O. dioica* and vertebrates. As in other species, the preferred location of the TATA box was 28–31 bp upstream.

We analysed profiles of methylated DNA enrichment around promoters using meDIP-chip (methylated DNA immunoprecipitation followed by chip) data [21] from ovaries, testes and tailbud stage embryos. Interestingly, we found that TATA-dependent sharp promoters had a higher average enrichment of DNA methylation in downstream gene bodies than TATA-less sharp promoters in embryos, ovaries and testes (Figure 4B). This trend was not explained by expression level (Figure S7A) or proximity to the promoter (Figure S7B) but did correspond to a higher frequency of CpGs (Figure S7C). A regression analysis showed that despite accounting for expression level (B = 0.06, *p* = 2.04 × 10^−8^), promoter width (B = −0.04, *p* = 2.47 × 10^−4^) and downstream CpG content (B = 0.26, *p* < 2 × 10^−16^) the presence of the most common core TATAA motif was a significant, independent, positive predictor (B = 0.27, *p* = 1.32 × 10^−12^) of downstream DNA-methylation levels in ovary, testis and tailbud, (the stage of development was not a significant predictor, overall fit of the model, *R*^2^ = 0.08). H3K4me3, which inhibits the interaction of DNA methyltransferases with histone proteins [31], was depleted (as was H3K4me2) at the TSS and in the downstream regions of TATA-dependent promoters, compared to TATA-less sharp promoters, in both the ovary and testis (Figure 4C). Together our findings reveal a specific association of gene body DNA methylation, and H3K4me3 depletion, with a TATA-dependent mode of TSS selection.

## Discussion

Here, we mapped sites of transcription initiation genome-wide at single nucleotide resolution across the life cycle of a marine chordate belonging to the sister group to vertebrates. Our data revealed a suite of of TSS-selection criteria in *O. dioica* (Figure 5) with features that are both shared with vertebrates and markedly different, particularly among maternal and spermatogenesis promoters.

**Figure 5:**
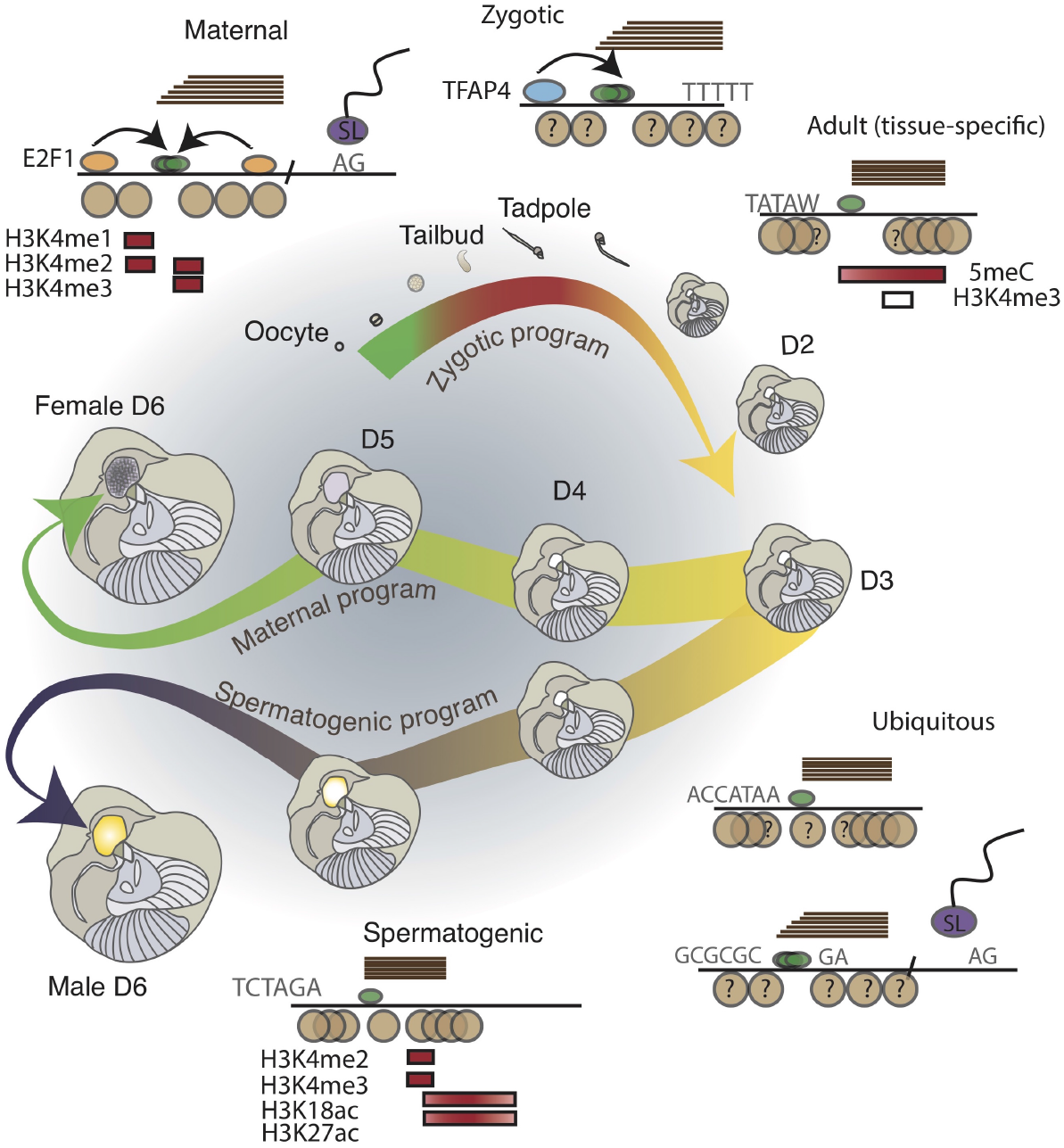
Promoter types in *O. dioica*. Schematic shows representative features of maternal, zygotic, ubiquitous (and ribosomal protein), tissue-specific and spermatogenic promoters with the key stages of the *O. dioica* life cycle. Horizontal lines represent positions of CAGE tags to indicate broad (staggered) or sharp (end-aligned) promoter architectures; green ovals represent pre-initiation complex sub-units. Motifs and dinucleotide enrichments are indicated. Brown circles represent positions of nucleosomes: no overlap indicates ordered positioning; question marks indicate lack of data. *Trans*-splicing with the spliced-leader (SL) at an AG acceptor site is also shown at promoter types where this is common. Boxes represent enrichments of histone modifications across maternal, spermatogenesis and adult promoters: red = enrichment; white = depletion.

A recent study in zebrafish found that maternal promoters are sharp (or multiple sharp) with a TATA-like upstream element whereas zygotic promoters are broad with transcription initiation guided by ordered nucleosome positioning [4]. In comparison, we found that in *O. dioica*, both maternal and zygotic promoters are broad. Moreover, we found evidence of nucleosome positioning as well as an enrichment for the binding of E2F1 at broad maternal promoters. These differences in maternal TSS-selection between zebrafish and *O. dioica* may be due to different modes of oogenesis and different sources of maternal transcripts. In zebrafish maternal transcripts originate from oocyte nuclei whereas in *O. dioica* the majority of maternal transcripts originate from terminally differentiated polyploid nurse nuclei within the single-cell coenocyst and are transported to oocytes through ring canals [32, 33].

Most maternal transcripts are *trans*-spliced in *O. dioica* [20] and this may have influenced the evolution of maternal promoter architectures. Since *trans*-splicing removes the 5’ end of a pre-mRNA (the “outron”) it follows that this sequence has little, if any, role in the post-transcriptional regulation of its mRNA. Indeed, one hypothesis for the function of *trans*-splicing in monocistrons is that it removes deleterious sequences at the 5’ end of an mRNA (e.g. premature start codons). There is mounting evidence that the SL sequence itself plays an important role in translational control, particularly for TOP mRNAs, which are *trans*-spliced in *O. dioica* [20]. We have shown here that the conserved TCT initiator sequence, which constitutes the first two nucleotides of the TOP motif, is absent at *O. dioica* ribosomal protein TSSs. We hypothesize that there is no requirement for a strict site of transcription initiation for *trans*-spliced genes since the spliced leader provides any necessary 5’ post-transcriptional regulatory motifs. Promoters of trans-spliced genes are then permitted to adopt a broad architecture governed by chromatin state rather than sequence motifs.

Our data also revealed a remarkable genome-wide shift in mode of TSS-selection during spermatogenesis to one associated with a position-specific core promoter motif (TCTAGA). One consequence of this shift is the developmental regulation of *trans*-splicing during spermatogenesis: many mRNAs that are *trans*-spliced in other stages (often in operons) are transcribed during spermatogenesis from an alternative TSS, driven by a TCTAGA promoter located downstream of the *trans*-splice acceptor site. This *trans*-splice acceptor site is thereby excluded from resulting mRNAs, which are no longer *trans*-spliced with the SL sequence. This may lead to a switch in the translational control of these transcripts to one that is independent of nutrient levels [20]. We hypothesise that this translational control is not required during the non-vitellogenic process of spermatogenesis. Nutrient-dependent control over initiation of meiosis has, however, been described in both sexes of *O. dioica* [34]. The TCTAGA promoter motif may play a role in this regulation in males if its binding by a transcription factor is nutrient-dependent.

A recent study found that genes with transcription-associated gene body methylation encode more highly conserved proteins with typical “housekeeping” functions [7]. We discovered a strong association of gene body DNA methylation with TATA-dependent promoters in *O. dioica*. This relationship is present during early development as well as in both the male and female germ lines, despite these differing substantially in their chromatin landscapes [21]. Promoters with the male-specific TCTAGA motif did not exhibit this downstream DNA methylation enrichment, despite this motif being position-specific and located in the expected TATA-box position. This indicates that gene body methylation in a subset of *O. dioica* genes is driven by core promoter features, specifically the TATA-box. A study in *C. intestinalis* found that gene bodies in near identical sets of genes are methylated in different cellular contexts [35], which is similar to our observations in *O. dioica*. This study also showed, however, that features within two ubiquitously expressed promoters are not the primary determinant of gene body DNA methylation. Analysis of additional *C. intestinalis* promoters may nevertheless reveal a relationship with the TATA-box similar to what we observe in *O. dioica*. Further exploration of sequence context in both species may also reveal a role for additional factors.

Given that DNA methylation in gene bodies suppresses transcription from alternative downstream promoters [9, 10] it is tempting to speculate that TATA-dependent sharp promoters employ DNA methylation as additional insurance for the strict positioning of transcription initiation. We also observed a depletion of H3K4me3 at, and downstream of, TATA-dependent promoters, in line with the inhibitory effect of H3K4me3 on DNA methyltransferases. Since TFIID is anchored at H3K4me3 on the +1 nucleosome [36] this indicates that TATA-dependent promoters are bound by TBP as part of an alternative complex. In yeast, TATA-dependent promoters are depleted of both TFIID and a nucleosome positioned downstream of the TSS and TBP is instead directed to the TATA-box by the SAGA complex [37]. Further investigation is required to establish whether or not a similar situation exists in metazoans.

Our results support previous findings of overlapping promoter codes [4], while revealing additional diversity and differential usage during complex developmental transitions. We provide the first links between acquisition of *trans*-splicing and the reorganization of promoter architectures for a conserved set of core metabolic genes, probably arising at least in part, because of regulatory sequences encoded in the SL. We also show shifts in TSS selection associated with a previously unidentified core promoter motif during the spermatogenic program. Further work on a range of additional models would provide a better framework in understanding the evolution of core promoter architectures, particularly with respect to innovations within major lineages.

## Methods and Materials

### Modified cap analysis of gene expression (CAGE)

Total RNA from each stage of development was isolated using RNAqueous Micro (Ambion) and treated by Terminator™ 5’-Phosphate-Dependent Exonuclease to deplete excess small RNAs. A modified CAGEscan protocol [11] was carried out at DNAFORM, Yokohama City, Japan. The standard CAGEscan protocol was modified in order to separate *trans*-spliced from non*-trans*-spliced transcripts by first using a custom designed 5’ linker, specific to the 5’ spliced leader sequence, before using standard linkers for non-*trans*-spliced mRNAs. Sequenced libraries for each stage therefore included only non-*trans*-spliced transcripts.

### Mapping reads

Illumina sequencing generated a total of 39,124,333 reads, 37 nt in length. We mapped these to the *O. dioica* reference genome [16] using Bowtie [38] with default parameters (allowing 2 mismatches per read). The 5’ coordinates of all uniquely mapping read (CAGE tag) alignments were extracted from the Bowtie output to give positions of CAGE transcription start sites (CTSSes), and the number of tags at each position was computed to give a tag count for each CTSS. We normalized tag counts to tags per million reads (tpm).

### Promoter types

We used the R package “CAGEr” [23] to cluster CTSSes into CAGE tag clusters, excluding those with < 1 tpm and singletons < 5 tpm, using a maximum distance of 20 bp between CTSSes within a cluster. We calculated the interquantile range (*q*_0.1_ − *q*_0.9_) of promoter widths (a measure of how broad/dispersed or peaked/focused a promoter’s TSS usage is that is more robust to expression level than using the full promoter width). We used this to group promoters into four classes using the mean and upper and lower quartiles as thresholds. We defined the upper and lower quartiles as “broad” and “sharp” respectively. We categorized promoters by CpG frequency in a 200 bp window centered on the dominant CTSS. Promoters with a CpG frequency in the upper quartile of CpG frequencies were classed as high CpG (HCG) and promoters with a CpG frequency in the lower quartile classed as low CpG (LCG). Using CAGEr we grouped all tag clusters with > 5 tpm across all stages into consensus promoter regions, using the interquantile range (*q*_0.1_ − *q*_0.9_) of tag cluster widths and a distance of 100 bp to merge clusters into one region. We used SOM clustering both at the level of individual CTSSes and consensus promoter regions to generate 25 expression profiles in each case.

### Shifting promoters

We calculated a shifting score and *p* of Kolmogorov-Smirnov test for all consensus promoters for all pairwise comparisons. We used a score >0.6 and FDR <0.01 to define a promoter shift – identifying promoters that have at least 60% of transcription initiation in the sample with lower expression occurring either upstream or downstream of transcription initiation in the compared sample.

### Assigning CTSSes to gene models and operons

We used Genoscope gene model predictions and annotations of polycistrons (http://www.genoscope.fr) to classify genes into operons and non-operons. A CTSS was associated with a gene model if it overlapped a gene body or its 500 bp upstream region. Using previously published CAGE data for *trans*-spliced transcripts [20], we classed a gene as SL *trans*-spliced if there was a SL CTSS within the gene body or within a 500 bp upstream region, if it was supported by > 1 tag count and if it had an ‘AG’ acceptor site motif immediately upstream.

### GO analysis

We used *O. dioica* GO annotations [22]. We used the Bioconductor GOstats package in R to compute hypergeometric *p*s for over-representation of GO terms in different sets of genes.

### Motif analyses

Over-represented motifs in core promoter regions were identified using MEME with default parameters on sequences in a 200 bp window, centred on the dominant CTSS within each tag cluster, for groups of CTSSes of interest. We also identified position-specific motifs (including initiator trinucleotides) by scanning core promoter regions for the occurrence of all possible k-mers (for k=1-6). We used TOMTOM to match position-weight matrices of motifs identified by MEME to known transcription factor binding sites [39]. We plotted the dinucleotide content of promoters using the R package “seqPattern”. We searched for TATA elements in the region 37–22 bp upstream of the dominant CTSS in each TC. We searched for TCTAGA motifs in the 22-52 bp and 52-101 bp upstream regions. We searched for ACCATAA motifs in the 32-72 bp upstream region. We used zygotic (CTSS SOM cluster 0_0) promoter sequences (200 bp centred on the dominant CTSS) from the tailbud stage to identify over-represented zygotic motifs. We used the “Biostrings” R package to scan (using a minimum score of 85%) the 101 bp upstream region with the position weight matrix discovered by MEME that matched the binding site for human TFAP4.

### ChIP-chip analysis

We analysed previously published meDIP-chip data and ChIP-chip data for E2F1, H3 and histone modifications H3K4me1, H3K4me2, H3K4me3 and H3K27me3 from mature *O. dioica* ovaries and testes [21]. We used the Bioconductor R package Ringo [40] for pre-processing all ChIP-chip data. Briefly, we normalized raw probe intensities from each sample (Cy5 channel) to corresponding input DNA probe intensities (Cy3 channel) by computing log_2_(Cy5/Cy3). We used the NimbleGen normalization method, which adjusts for systematic dye and labeling biases by subtracting from individual log2 ratios the Tukey’s biweight mean, computed across each sample’s log_2_ ratios. To reduce noise in the data we smoothened the normalized log_2_ ratios using a running median across a 150 bp window (the approximate size of a single nucleosome) with a minimum threshold of 3 non-zero probes. For each group of promoters we plotted the mean log_2_ ratio at each probe position for all probes in a 1000 bp window centred on the dominant CTSSes of promoter regions of interest. We excluded promoters with flanking regions that overlap. We defined regions of ChIP-enrichment genome-wide as previously described [21].

### Operon transcription analysis

We used tiling array data generated from *O. dioica* testes and ovaries [22] to categorize genes within operons as expressed or silent. We then defined an operon as expressed if any of its genes are classed as expressed. We intersected H3K4me3 ChIP-enriched regions with operon promoters, as well as potential promoters of internal operon genes, using the region 500 bp upstream and 100 bp downstream of annotated start sites. Any overlap was defined as a presence of H3K4me3 in a candidate promoter region.

### 5’ RACE

RACE was performed using SMARTER RACE kit from Clontech according to manual.

### Re-analysis of mouse testis CAGE data

Analysis of TSS data followed that found in [23] using data downloaded from http://promshift.genereg.net/CAGEr/InputData/ consisting of TSSs from 8 stages of mouse testis development. Briefly, we used the CAGEr [23] package to normalize tag counts and cluster TSSs into TCs for each stage. We plotted the frequency of TCTAGA motif around TSSs from each stage, sorted by the width of TCs and saw no enrichment. We then used a self organizing map (SOM) to cluster the expression profiles of each TSS and identified a cluster with expression specific to later development which was previously annotated as being enriched for TSSs associated with spermatogenesis genes [23]. We plotted the TCTAGA frequency around the TSSs of this cluster and also saw no enrichment.

### Search for TCTAGA and TATA-box motifs in *C. intestinalis* promoters

We searched the 100 bp region upstream of all annotated Ensembl 87 KH *C. intestinalis* genes for “TCTAGA” motif and the consensus TATA-box motif “TATAWAR” using the “Biostrings” R package.

## Acknowledgments

We thank Matthias Harbers and his staff at DNAFORM (Yokohama, Japan) for their assistance in the development of the modified CAGE protocol. We thank Jean-Marie Bouquet, Magnus Reeve and Anne Aasjord for supplying animals as part of the animal culture facility. The authors declare that they have no competing interests.

The datasets generated and/or analysed during the current study are available in the NCBI Gene Expression Omnibus under accession number GSE78794 and GSE78915

